# Phylogenomics and barcoding of *Panax*: toward the identification of ginseng species

**DOI:** 10.1101/244780

**Authors:** V. Manzanilla, A. Kool, Nhat L. Nguyen, H. Nong Van, H. Le Thi Thu, H.J. de Boer

**Affiliations:** The Natural History Museum, University of Oslo, Oslo, Norway; Institute of Genome Research, Vietnam Academy of Science and Technology, 18 Hoang Quoc Viet, Cau Giay, Hanoi, Vietnam

**Keywords:** barcoding, genome, ginseng, marker, mPTP, NGS, Panax, Phylogenomics, Plastid

## Abstract

**Background:** The economic value of ginseng in the global medicinal plant trade is estimated to be in excess of US$2.1 billion. At the same time, the evolutionary placement of ginseng (*Panax ginseng*) and the complex evolutionary history of the genus is poorly understood despite several molecular phylogenetic studies. In this study, we use a full plastome phylogenomic framework to resolve relationships in *Panax* and to identify molecular markers for species discrimination.

**Results:** We used high-throughput sequencing of MBD2-Fc fractionated *Panax* DNA to supplement publicly available plastid genomes to create a phylogeny based on fully assembled and annotated plastid genomes from 60 accessions of 8 species. The plastome phylogeny based on a 163 kbp matrix resolves the sister relationship of *Panax ginseng* with *P. quinquefolius*. The closely related species *P. vietnamensis* is supported as sister of *P. japonicus*. The plastome matrix also shows that the markers *trnC-rps16, trnS-trnG*, and *trnE-trnM* could be used for unambiguous molecular identification of all the represented species in the genus.

**Conclusions:** MBD2 depletion reduces the cost of plastome sequencing, which makes it a cost-effective alternative to Sanger sequencing based DNA barcoding for molecular identification. The plastome phylogeny provides a robust framework that can be used to study the evolution of morphological characters and biosynthesis pathways of ginsengosides for phylogenetic bioprospecting. Molecular identification of ginseng species is essential for authenticating ginseng in international trade and it provides an incentive for manufacturers to create authentic products with verified ingredients.

## Background

Ginseng has been used in traditional medicine in China for thousands of years [1], but it was not until early 18th century that long-term, intense harvest nearly extirpated *Panax ginseng* C.A.Mey. from the wild [2]. Demand for ginseng roots in the 18th century also fuelled a subsequent boom in wild-harvesting American ginseng (*P. quinquefolius* L.) that decimated wild populations in North America [3]. Today wild *P. ginseng* occurs in only a few localities in Russia and China, with the largest distribution in the southern part of the Sikhote-Alin mountain range [4]. *P. ginseng* is Red-Listed in Russia, and roots and parts thereof from Russian populations are CITES Appendix II/NC listed [5]. Many other Asian ginseng species are also endangered but preliminary data is only available for wild-harvesting and conservation of *P. assamicus* R.N.Banerjee (synonym of *P. bipinnatifidus* var. *angustifolius* (Burkill) J.Wen) [6], *P. japonicus* (T.Nees) C.A.Mey. [7] and *P. pseudoginseng* Wall. [8, 9].

Elucidating the evolutionary relationships among species in the genus is essential to understand evolution of this Holarctic disjunct genus, but also evolution of derived secondary metabolite pathways. In addition, a phylogenetic framework can be used to develop accurate molecular identification of *Panax*, and enable identification of ginseng material in trade, both crude drugs and derived products, which is essential for conservation efforts and protection of the remaining wild populations of *P. ginseng* and related *Panax* species, since all may be under the pressure of illegal harvesting and international trade [10]. Furthermore, identification of *Panax* species and authentication of derived products is of great commercial importance as authentic ginseng is costly and the incentive for substitution is significant.

The phylogeny of *Panax* has been studied using several molecular markers, but lack of variation in the most commonly used markers highlight an important limitation of the method. The nuclear ribosomal ITS yields insufficient resolution for accurate species assignment [19] and even using multiple markers in combination, *matK, trnD, psbK-psbI, rbcL* and *ycf1* have a limited accuracy in identification of *Panax* species [20, 21]. The mutation rate of the studied markers does not allow a fine scale resolution, and is insufficient for identification of all *Panax* species and cultivars. The question of what species are in trade remains a mystery. Aside from phylogenetic approaches, a multitude of molecular and chemical analysis approaches have been developed and applied, including Arbitrarily Primed Polymerase Chain Reaction (AP-PCR) [11], PCR-Restriction Fragment Length Polymorphism (PCR-RFLP) and Mutant Allele Specific Amplification (MASA) [12], Random Amplified Polymorphic DNA (RAPD) and High Performance Liquid Chromatography [13], Fourier Transformed-Infrared Spectroscopy (FT-IR) [14], Two-Dimensional Correlation Infrared Spectroscopy (2D-IR) [14], Multiplex Amplification Refractory Mutation System-PCR (MARMS) [15, 16], Microchip Electrophoresis Laser-Induced Fluorescence Detection [17], and microsatellite markers [18]. Most methods have focused on either positive identification of *P. ginseng*, or distinguishing *P. ginseng* and *P. quinquefolius* L., but most have limited resolution in detecting infraspecific or interspecific substitution, especially with poorly known congeneric species.

Suitability of molecular markers is often measured in interspecific distance using distance methods to estimate the number of variable sites or pairwise distances between sequences. Most current methods are based on the Refined Single Linkage (RESL) algorithm implemented in BOLD [22] or clustering on distance matrices (Crop [23], OBITools [24], UCLUST [25], and Vsearch [26]) and ideally set a threshold to distinguish between intraspecific and interspecific variation, sometimes referred to as the “barcoding gap” [27]. Several programs and software packages determine and visualize barcoding gaps, including Automatic Barcode Gap Discovery (ABGD) [28] and Spider [29]. These distance-based methods are fast and suitable for large datasets, but they are not always biologically meaningful, especially when the species groups have complex evolutionary histories, including incomplete lineage sorting, and hybridization [30, 31]. As an alternative, tree-based methods offer several advantages compared to distance based methods. First, these methods do not work with a specified threshold (% variation, no barcoding gap) and second, these accommodate evolutionary processes, making them particularly suitable for species delimitation and identification. Several studies have shown that these methods are also more sensitive and more powerful for accurate species discrimination [32]. Recently proposed methods include the Generalized Mixed Yule Coalescent (GMYC) [33], Bayesian species identification using the multispecies coalescent (MSC) model [34], and Poisson Tree Processes (PTP, mPTP) [25, 32]. Despite constant methodological improvements, there is no silver bullet for species delimitation and concerns have been raised that species delimitation approaches are sensitive to the structure of the data tested [35]. Species delimitation methods assess speciation and coalescent processes but also the data structure of the selected markers [35]. From a marker development perspective, tree based methods provide an opportunity to increase the quality of the selection process of the barcoding markers. Here we use the mPTP approach [32] to test if speciation processes are supported by the barcoding markers and accordingly choose the best markers for delimitation of *Panax* species. mPTP method has the advantage of being computationally efficient, while at the same time accommodating better to population-specific and sampling characteristics than PTP and GYMC [32].

## Evolution and phylogenetics of *Panax*

Phylogenetic studies of the Araliaceae based on nuclear ribosomal DNA and chloroplast DNA have circumscribed four major monophyletic groups (the Asian Palmate group, the Polyscias-Pseudopanax group, the Aralia-Panax group, and the greater Raukaua group) [36, 37]. However deep nodes are not well-supported to date [36, 37], and a broad sampling within Aralioideae is necessary to obtain an accurate placement of the Aralia-Panax group. Monophyly of the genus *Panax* (Araliaceae) is well supported by morphological synapomorphies, such as palmately compound leaves, a whorled leaf arrangement, a single terminal inflorescence, valvate petals in floral buds, and a bi- or tricarpellate ovary, as well as by several molecular phylogenies [20, 38]. A number of species have emerged from the complex of subspecies of *P. pseudoginseng* in the 1970s, and taxonomic studies have resulted in the description of various new species [38–40]. Currently thirteen species of ginseng are recognized with broad consensus [38, 41], but publication of new taxa at species, subspecies and variety level are common [42, 43].

Previous molecular phylogenies support *P. stipuleatus* H.T.Tsai & K.M.Feng and *P. trifolius* L. as the sister group of all other ginseng species. Nevertheless the placement of several other species still remains unclear (e.g., *P. binnatifidus, P. ginseng, P. japonicus, P. quiquefolius, P. vietnamensis* Ha & Grushv., *P. wangianus* S.C.Sun, *P. zingiberensis* C.Y.Wu & Feng). Species delimitation within the genus is problematic due to species of tetraploid origin (e.g., *P. bipinnatifidus, P. ginseng, P. japonicus*, and *P. quinquefolius* [44]), recent speciation events [20], high intraspecific morphological variation (e.g., *P. pseudoginseng* Wall.) and ancient genome duplication events [41, 45].

Phylogenetic studies have explored evolutionary relationships in Araliaceae with standard phylogenetic markers, such as the nuclear ribosomal ITS [19, 36, 38, 41, 44–46] and several plastid markers [19–21, 41]. More recently, an attempt with seven nuclear genes was tested with moderate results (*PGN7, W8, W28, Z7, Z14, Z15, Z16*) [20]. The topologies obtained were conflicting and non-consistent with previous evolutionary inferences of the genus, which is likely a result of multiple copies of nuclear genes and ancient whole genome duplication events [47]. Whole genome data have also been used to design microsatellites for species identification, but these have found limited application [18, 48–52]. Extensive population genetic studies have been done only on *P. quinquefolius* [53–59] and *P. ginseng* [60, 61] due to their major economic importance.

Developments in high throughput sequencing have provided new approaches for genome sequencing: increasing outputs and decreasing costs have made this a cost-effective alternative to Sanger-based amplicon sequencing [62, 63]. Full plastid genome sequencing, *i. e*. plastome sequencing, has been proposed as an augmented approach to DNA barcoding [64, 65], and is a straightforward method that recovers all standard barcodes plus the full plastome. The limited costs of shotgun sequencing and the availability of a number of Araliaceae reference plastomes facilitates the study of relationships in the family. Plastome phylogenies have helped disentangle evolutionary relationship in a number of plant clades [66], including Poales [67], magnoliids [68], *Pinus* [69], *Amborella* [67], *Equisetum* [70], and *Camellia* [71]. Single-copy nuclear genes have corroborated the robustness of plastome phylogenies [72–75], however plastome phylogenies reflect only maternal inheritance, and as such will not always be representative species trees. An advantage of plastome data for phylogenetic studies is the low mutation rate of plastid sequences, the abundance of plastid DNA in most material [76] and the low cost of generating whole plastid genomes with high throughput sequencing.

In total DNA, the proportion of plastid DNA typically constitutes only ~0.01–13% depending on the size of the nuclear genome, tissue and season [77–79]. Shot-gun sequencing studies might have relatively low efficacy in plastid genome recovery due to the small proportion of plastid DNA in the total DNA. Ginseng species have a large genome size of 5–10 Gb [80, 81], and one can expect a proportion of plastid DNA of 1–5 % in the gDNA [79], which makes shotgun sequencing relatively ineffective in obtaining full plastome data. Several methods have been developed for enriching plastid content prior to sequencing (for a discussion see Du et *al*. [82]. We apply a new plastid enrichment method to improve the shotgun sequencing efficacy, that utilizes the low methylation of the plastid genome compared to the nuclear genome [83]. The method uses the methyl-CpG-binding domain (MBD2) to partition fragments of genomic DNA into a methylation-poor fraction (e.g. enriched for plastid) and a methylation-rich fraction (e.g. depleted in plastid) [84]. This method has the advantage that it uses a small quantity of dry material (below 40 mg) and is suitable for non-model organisms.

This study has four main aims: (1) to construct a well-supported phylogeny of the genus *Panax*, while testing if the full plastome data yield sufficient variation to support and resolve phylogenetic relations in *Panax*, and specifically the position of the economically important *P. ginseng*; (2) to test if MBD2 can be used to fractionate *Panax* DNA into eukaryotic nuclear (methyl-CpG-rich) vs. organellar (methyl-CpG-poor) elements, and subsequently sequence the MBD2 depleted DNA to optimize plastome read yield; (3) to determine if the plastid genome can be used for molecular identification of traded species; and 4) to make a case for the need of a resolved plastome phylogeny to be used to design short markers for *Panax* species identification from processed ginseng products.

## Materials and methods

### Sampling

Fresh material of three species, *P. bipinnatifidus, P. stipuleanatus*, and *P. vietnamensis* (*2*), was sampled in Vietnam (Table 1) and 57 selected Araliaceae plastid genomes from across the Araliaceae family were downloaded from open data repositories (Table S1) [20, 85–97]. At least two individuals or species were selected per genus, but for *Panax* we used 38 plastid genomes from eight species. *Hydrocotyle verticillata* was selected as outgroup based on its early divergence within Araliaceae [44].

**Table 1.**
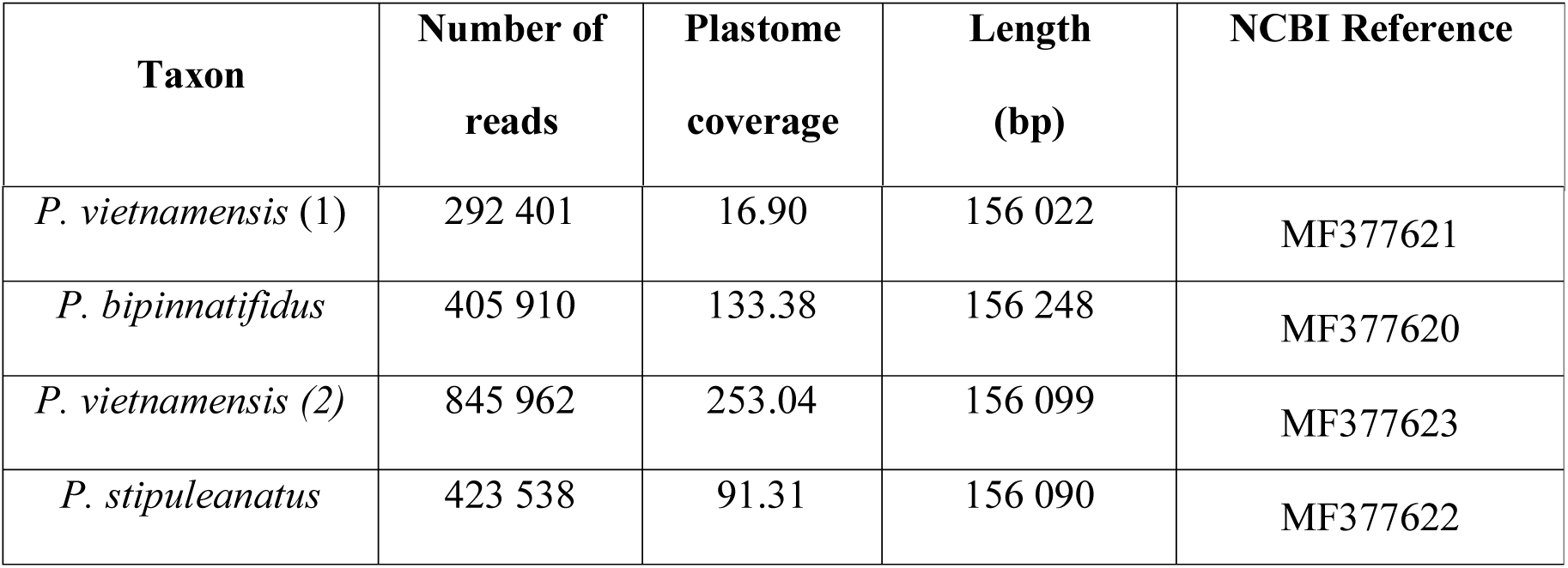
Summary information for the four assembled plastome genomes.

| Taxon | Number of reads | Plastome coverage | Length (bp) | NCBI Reference |
| --- | --- | --- | --- | --- |
| *P. vietnamensis* (1) | 292 401 | 16.90 | 156 022 | MF377621 |
| *P. bipinnatifidus* | 405 910 | 133.38 | 156 248 | MF377620 |
| *P. vietnamensis* (2) | 845 962 | 253.04 | 156 099 | MF377623 |
| *P. stipuleanatus* | 423 538 | 91.31 | 156 090 | MF377622 |

### Library preparation and sequencing

We extracted total DNA from two individuals of those sample collected in Vietnam, using a Qiagen DNeasy plant extraction kit with the provided protocol. The total DNA was quantified prior to library preparation to assess DNA quantity, fragmentation and fragment length distribution on a Fragment Analyzer (Advanced Analytical Technologies, Inc., Ankeny, USA) using the High Sensitivity genomic DNA Reagent Kit (50–40,000 bp) (Fig. S1). We selected one individual per extracted sample based on the yield and fragment size of the total DNA. The selected samples had average fragment sizes in excess of 10 kbp and a minimum DNA concentration of 4.77 ng/µl (Fig. S1).

We used a NEBNext Microbiome DNA Enrichment Kit (New England Biolabs, Ipswich, Massachusetts, USA) that uses IgG1 fused to the human methyl-CpG-binding domain (together “MBD2-Fc”) to pull down a methyl-CpG-enriched fraction from a bead-associated element, leaving a methyl-depleted fraction in the supernatant. About 400 ng template DNA extract was used per sample and the manufacturers recommendations were respected with the following exceptions. The non-methylated DNA fractions were purified using 0.9X AMpure XP beads (Beckman Coulter, Brea, CA, USA) and eluted in 40 µl 1X TE buffer. To capture the methylated DNA, we followed the manufacturer’s protocol. Quality control in terms of size, purity and molar concentration (nmol/l) of both the methylated and the non-methylated fractions were measured using a Fragment Analyzer (Advanced Analytical Technologies Inc., USA) with a DNF-488–33 HS dsDNA Reagent Kit. The DNA was subsequently sheared to ~400 bp fragments using a M220 Focused Ultrasonicator (Covaris Inc., Woburn, MA, USA) using microTUBES-50 (Covaris Inc.). We used the NEBNext Fast DNA Library Prep Set for Ion Torrent (NEB) for end repair and adapter ligation of the sheared DNA. The samples were indexed using the IonXpress Barcode Adapter kit (ThermoFischer, Waltham, MA, USA). For each of the four samples both fractions, methyl-CpG-enriched and methyl-CpG-depleted, were indexed and sequenced. After adapter ligation, the four methyl-CpG-enriched fractions were pooled in one library and the four methyl-CpG-depleted fractions were pooled in another library. The adapter-ligated libraries were size selected (450–540 bp) using a BluePippin (Sage Science, Beverly, MA, USA), and subsequently amplified using the NEBNext Fast DNA Library Prep Set for Ion Torrent kit using 12 PCR cycles. The amplified libraries were purified twice using 0.7X AMpure XP beads. The purified amplified libraries were loaded on the sequencing chips using an Ion Chef (LT) and sequenced on an Ion Torrent Personal Genome Machine (LT) using Ion 318 v2 chips (LT) and the Ion PGM Sequencing 400 kit (LT).

### Bioinformatic analyses and assembly

Sequencing reads were demultiplexed into FASTQ files using Flexbar version 3.0.3. Trimmomatic version 0.36 [98] was used for adapter trimming and quality filtering of reads using a sliding window of 15 bp and an average Phred threshold of 20. Low-end quality bases below a Phred score of 20 were removed, and only reads longer than 100 bp were retained. MITOBim version 1.7 [99] was used for assembly of the single-end Ion Torrent reads using iterative mapping with *in silico* baiting using the following reference plastomes, *P. vietnamensis* (KP036470) and *P. stipuleanatus* (KX247147).

Inverted repeats and ambiguous portions of the assembly were resequenced using Sanger sequencing. Specific primers were designed and used for DNA amplification of interest regions. PCR was performed on a Mastercycler^®^ Pro (Eppendorf, USA) in a 20 µl final volume containing 2.5 µM of each primer, 1 mM of each dNTP, 10X DreamTaq Buffer, 0.75 U DreamTaq DNA polymerase (ThermoFisher Scientific, USA) and deionized water. The PCR cycling conditions included a sample denaturation step at 94^o^C for 2 minutes followed by 35 cycles of denaturation at 94 °C for 30 seconds, primer annealing at 50–55 °C for 30 seconds and primer extension at 72 °C for 1 minute, followed by a final extension step at 72 °C for 5 minutes. PCR products were then purified using GeneJET PCR Purification Kit (ThermoFisher Scientific, USA). Sanger sequencing was performed on an ABI 3500 Genetic Analyzer system using BigDye Terminator v3.1 Cycle Sequencing Kit. Cycle sequencing was performed on a Veriti Thermal Cycler (Applied Biosystems, USA) using 3.2 µM of each primer, 200 ng purified PCR product, 5X BigDye Sequencing Buffer, 2.5X Ready Reaction Premix and deionized water in a 20 µl final volume. The thermocycling conditions included 1 minute at 96 °C followed by 25 cycles of denaturation at 96 °C for 1 minute, primer annealing at 50 °C for 5 seconds and primer extension at 60 °C for 4 minutes, followed by a holding step at 4 °C. Extension products were purified using ethanol/EDTA precipitation with 5 µl of EDTA 125 mM, 60 µl of absolute ethanol. Purified products were denatured at 95 °C for 5 minutes using 10 µl Hi-Di Formamide. DNA electrophoresis was performed in 80 cm × 50 µ capillary with POP-4 polymer (Applied Biosystems, USA).

In order to test the efficacy of the NEBNext Microbiome DNA Enrichment Kit the proportion of reads belonging to the plastome was estimated for both the methylated and the non-methylated fraction. The *P. ginseng* whole genome sequencing SRR19873 experiment was used to estimate the starting proportion of plastome reads, by mapping the reads against the plastid genome of *P. ginseng* (NC_006290) using Bowtie 2. Association of reads to their taxonomic identification and organelles, was made using a tailored database of *Panax* plastome data representing the same data as that downloaded from public repositories for the phylogenetic analyses. For the mitochondrial data, all angiosperm mitochondrion genomes available on NCBI were used, and for the microbiome all remaining reads were blasted against the full NCBI database. Taxonomic identifications were retrieved using the lowest common ancestor (LCP) algorithm in Megan version 5.11.3, with minimum read length of 150 bp and at least 10 reads for each taxon identified with an e-value of 1e-20 or less. The proportion of plastid DNA in the gDNA was estimated using Bowtie2 by mapping the proportion of reads belonging to the plastid genome for *P. ginseng* (following SRR experiment SRR1181600).

The plastid genomes were annotated using Geneious version 6.1, and annotations of exons and introns were manually checked by alignment with their respective genes in the same annotated species genome. Representative maps of the chloroplast genomes were created using OGDraw (Organellar Genome Draw, [100]).

## Phylogenomics

The matrix for phylogenomic analyses consisted of complete aligned plastid genomes, and the global alignment was done using MAFFT version 7.3 [101] with local re-alignment using MUSCLE version 3.8.31 [102], and manual adjustments where necessary. Aligned DNA sequences have been deposited in the Open Science Framework directory (https://osf.io/ryuz6). The final matrix has a total length of 163,499 bp for a total of 61 individuals with no missing data. Single nucleotide polymorphisms (SNPs) were visualized using Circos version 0.69 [103]. Relationships from the nucleotide matrix were inferred using Maximum Likelihood (ML) and Bayesian inference. First, an un-partitioned phylogenetic analysis was performed to estimate a single nucleotide substitution model and branch length parameters for all characters. Next, the data was partitioned in coding regions, introns and intergenic spacers, and a best-fit partitioning scheme for the combined dataset was determined using PartitionFinder version 2.1.1 [104] using the Bayesian Information Criterion. Branch lengths were linked across partitions.

The dataset was analyzed using RAxML version 8.2.10 [105] and mrBayes version 3.2.6 [106]. RAxML and Bayesian searches used the partition model determined by PartitionFinder. For the ML analyses, tree searches and bootstrapping were conducted simultaneously with 1000 bootstrap replicates. Bayesian analysis were started using a random starting tree and were run for a total of 10 million generations, sampling every 1000 generations. Four Markov runs were conducted with 8 chains per run. We used AWTY to assess the convergence of the analyses [107]. Conflicting data within ML and Bayesian analyses were visualized and explored using the R package phangorn using the *consensusNet* function [108].

## Barcoding - mPTP

Suitable barcoding markers were selected by extracting the SNP density over the plastid genome alignment of all *Panax* species and individuals included in this study (matrix available as supplementary data on OSF). We used SNP-sites version 2.3.2 [109] to extract the SNP positions from the alignment of a matrix containing only the *Panax* species, and created bins every 800 bp using Bedtools version 2.26.0 [110] (script available on OSF) and plotted the SNP density using Circos [103] (Fig. 1). The coordinates of each annotation on the aligned *Panax* species matrix were found using a reference consisting of the four annotated genomes produced in this study, and subsequently exported to Circos. We selected the most variable regions and designed suitable primers for these regions (Fig. 5). From the matrix used for the Aralioideae, we extracted 15 plastid markers (Fig. 5) and download ITS sequences for the *Aralia-Panax* group (Fig. 3 & 5) (Table S1). We performed maximum likelihood analyses on individual and concatenated matrices using RAxML. mPTP analyses were performed using the ML trees from the individual and concatenated markers, and using the MCMC algorithm with two chains and the Likelihood Ratio Test set to 0.01.

**Figure 1.**
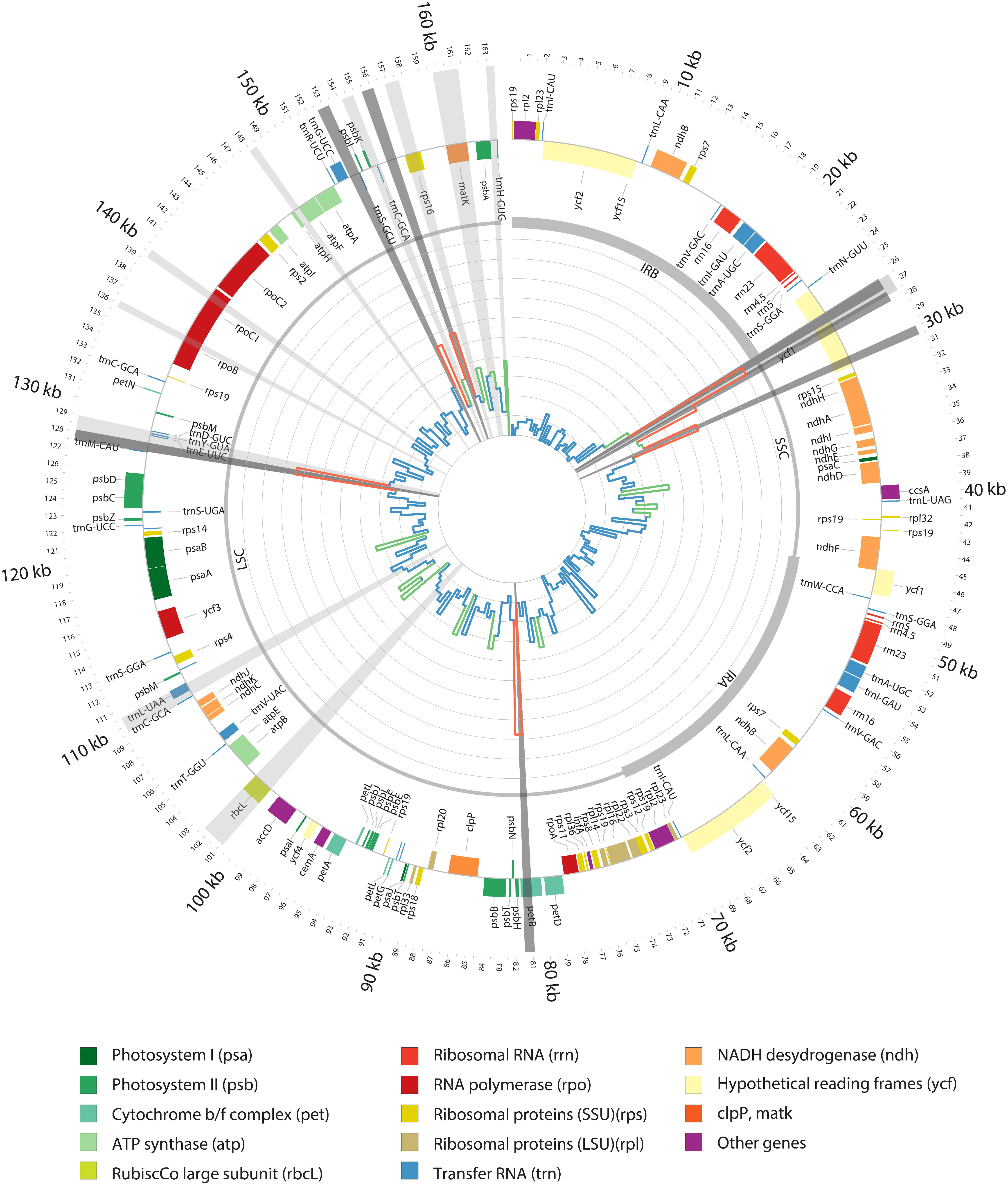
Plastid genome representation of the 38 aligned *Panax* genomes. The internal histogram plot represents the SNPs density over the alignment of the plastid genomes of *Panax* genus. The colours indicate when the standard deviation of the bin falls in different interval compare to the average standard deviation, between 0 and 1 in blue (low variation), between 1 to 2 in green (moderate variation) and over two in red (highly variable). Inverted repeats A and B (IRA and IRB), large single copy (LSC) and small single copy (SSC) are shown in the inner circle by different line weights. Genes shown outside the outer circle are transcribed clockwise, and those inside are transcribed counter clockwise. Genes belonging to different functional groups are color-coded. Radial grey highlights show the regions in focus of study, light grey previously used barcodes, in dark grey newly developed barcodes.

## Results

### Ion torrent sequencing

After filtering out low-quality reads, 1.9 out of 3.3 and 3.3 out of 4.9 million reads were retained for the pooled MDB2 depleted and enriched fractions respectively. The chloroplast assemblies covered the entire circular plastid genome for all four accessions for the MDB2 depleted fraction (Figs. S2, S3, S4, S5; Table 1). The Sanger generated plastid sequences confirmed the genome assemblies in 18 regions, and also confirmed sequences of the inverted repeat. Complete lengths of the four plastid genomes ranged from 156 036 bp to 156 302bp (Table 3). All four plastid genomes had the same genome structure and gene arrangement as that of the already assembled *Panax* plastid genomes.

### Methylation enrichment

The Fragment Analyzer results showed that DNA quantity and fragmentation differed for the four DNA samples (Fig. S1), and the results were used to normalize concentrations for subsequent capture. DNA concentrations after capture and fragment size selection are much lower for the methyl-depleted fraction compare to the methyl-CpG-enriched fraction (Fig. 2). The success of the fragment size selection was relatively poor for one of the *P. vietnamensis*. and resulted in a poorer quality in the sequencing and enrichment due to the excessive abundance of short DNA fragments. The shorter reads for *P. vietnamensis*. yielded a lower coverage for its genome assembly (16.9 X) (Table 1).

**Figure 2.**
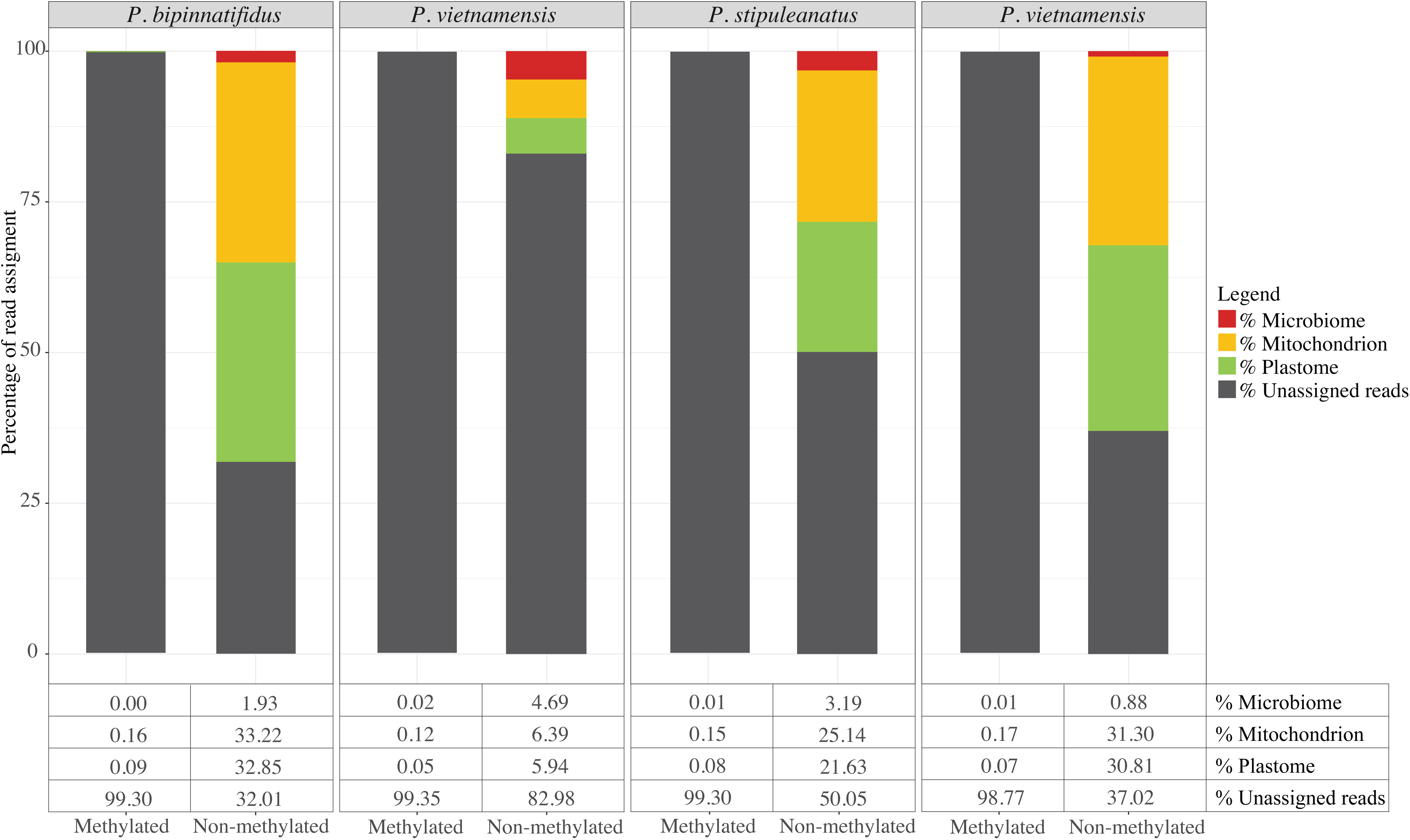
Proportion of mitochondrial, plastid and microbiome reads from the methylated and non-methylated fraction for the four enriched libraries. These results show that the enrichment procedure successfully capture more than 99.5% of the plastid sequences contained in the total DNA.

The enrichment and depletion of methylated DNA by pulling down a methyl-CpG-enriched fraction and leaving a methyl-depleted fraction drastically increased the proportion of organellar DNA within the depleted fraction. *P. ginseng* SRR experimental data had 5.63% plastid genome reads. In the methylation-depleted fraction, we found a variation of plastome reads ranging from 6% to 33%. In the methylation-enriched fraction, less than 1% of the reads are from the plastome. The enrichment also increased microbiome contamination in the depleted fraction from 0.8% to 4%. Overall, one of the *P. vietnamensis* samples was the least successful sample in the enrichment and yielded fewer and shorter reads.

### Phylogenetic analyses

Alignment of the plastid genomes for phylogenetic analyses were consistent in length throughout the dataset. Based on the alignment, average plastome pairwise identity for the Araliaceae family is 83% and 99.2% for the *Panax* clade. The percentage of identical sites is 83.9% and 96.8% respectively. The global plastome alignment has a matrix length of 163 499 bp. Coding regions, introns and intergenic spacers represented 259 original partition schemes, and the best-fit partitioning scheme from PartitionFinder divided the data into 73 partitions (Table 3).

Inspection of the posterior probabilities calculated using AWTY, yielded an estimated burnin of 10% for the Bayesian analysis. Phylogenetic analyses revealed significant divergence between major clades of the Araliaceae family. The ML and Bayesian trees showed strongly supported clades for all genera of the family (Fig. 3). Furthermore, the tree shows maximum support for each species of *Panax* included in the analyses. All intergeneric and infrageneric relationships were strongly supported (Fig. 3).

**Figure 3.**
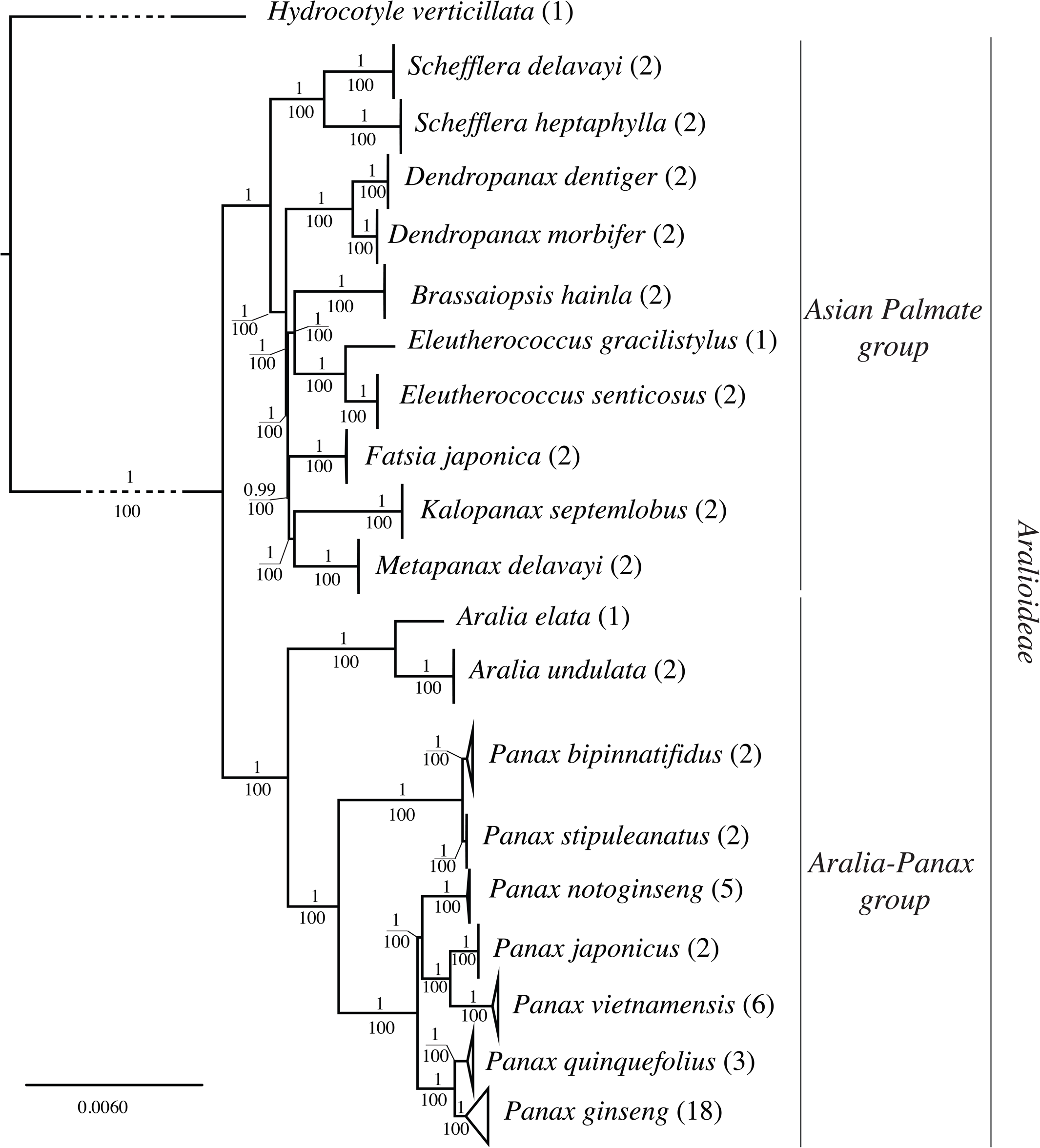
Bayesian tree inferred from the 61 plastid genomes from the Aralioideae tribe. Numbers above branches indicate posterior probabilities and below the ML bootstrap values.

The basal node segregates two clades, one clade includes two genera, *Aralia* and *Panax*. The second clade includes *Schefflera, Fatsia, Eleutherococcus, Kalopanax, Metapanax, Brassaiopsis*, and *Dendropanax*. All species included in the study are monophyletic and have maximum support in both Bayesian and ML analyses.

#### The Araliaceae clade

The Araliaceae clade showed maximum support in the phylogeny except for the *Fatsia* clade, where the support is 99.6%. *Schefflera* is sister to the rest of the clade, followed by *Dendropanax*, then a clade with *Brassaiopsis/Eleutherococcus* and finally a clade with *Fatsia*/*Kalopanax*/*Metapanax*. A comparison of the partitioned and non-partitioned analyses shows no differences in topology and support in the *Aralia-Panax* clade, but does in the remaining Araliaceae clade.

#### The *Aralia-Panax* clade

The genus *Panax* is monophyletic and *Aralia*, represented by two species, *A. elata* and *A. undulata*, is the sister group to the genus *Panax. Panax stipuleatus* and *P. binnatifidus* form a distinct clade sister to a clade consisting of *P. notoginseng* and its sister group of *P. vietnamensis* and *P. japonicus*, which as a whole is sister to *P. quinquefolius* and *P. ginseng*.

The consensus network was computed from the two Bayesian runs after discarding 10% burnin (Fig. 4). The network analysis shows two main conflicts in the data, one within the *P. ginseng* clade and another within the *P. vietnamensis* clade. Both clades have very little intraspecific variation (soft incongruence), and more variable markers are needed to segregate the different individuals correctly for these two species.

**Figure 4.**
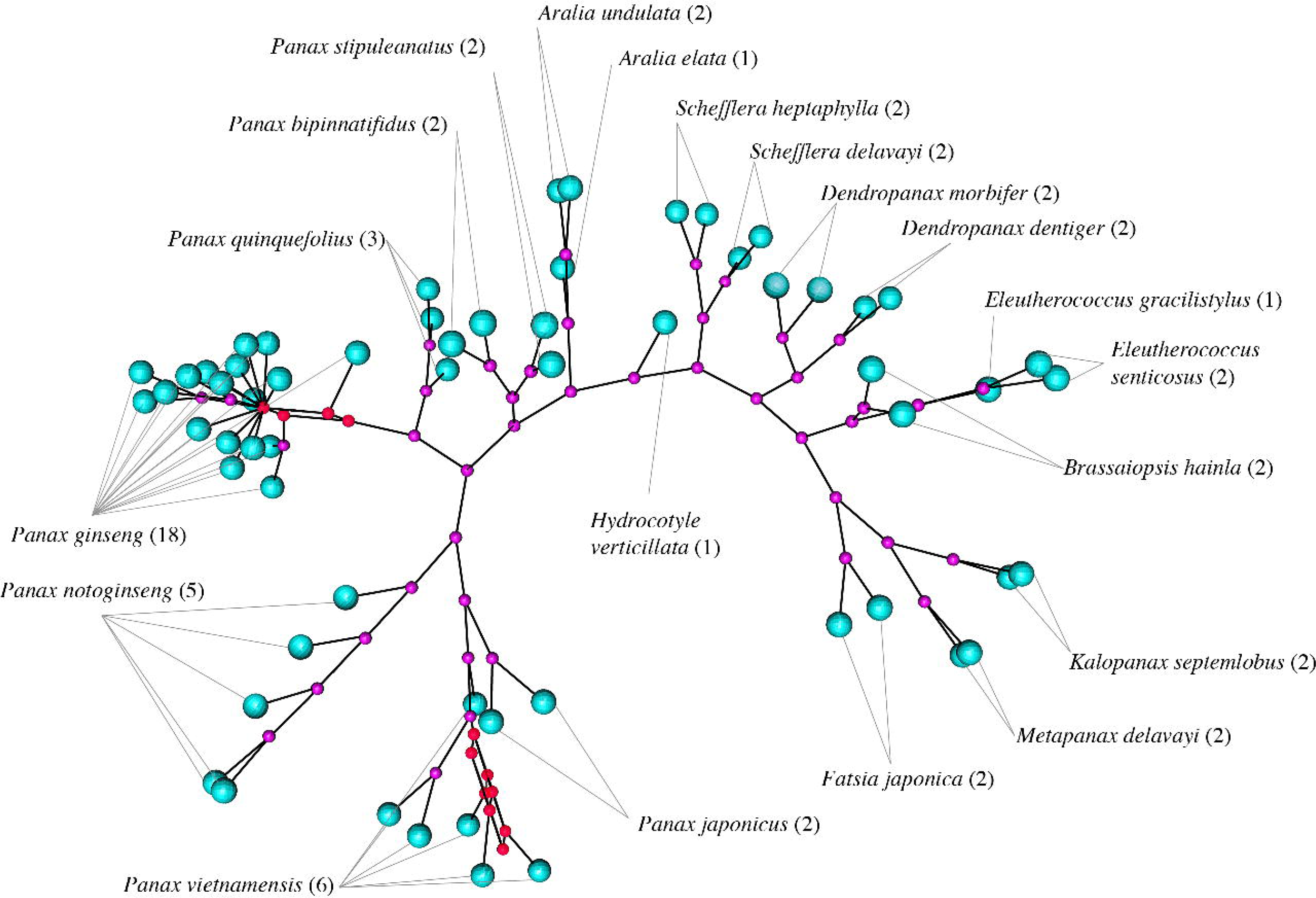
Consensus network from Bayesian runs with 10% burn in. This network shows two main conflicts in the data, one on the *P. ginseng* clade, where there is very little intraspecific variation and one with the clade of *P. vietnamensis*. The nodes are represented in pink and the tips in blue. The red nodes show the two splits within the data.

### Barcoding analyses

The SNP density analyses retrieved 2,052 SNPs over the full plastid alignment. We identified three regions (Figs. 1, 5) that are suitable barcoding markers. Each of these regions has on average of 83 SNPs within *Panax* (Fig. 5). Individual marker phylogenies of these regions are suitable to segregate most of the species clades (supplementary material S7-S11). The exceptions are the two sister pairs, *P. quinquefolius* and *P. ginseng*, and *P. binnatifidus* and *P. stipuleatus*, where the bootstrap supports are weaker, leading to inference of single clades. The ML phylogeny of the concatenated markers, fully supports all species clades, except *P. binnatifidus* and *P. stipuleatus* (Figs. 5, S14).

**Figure 5.**
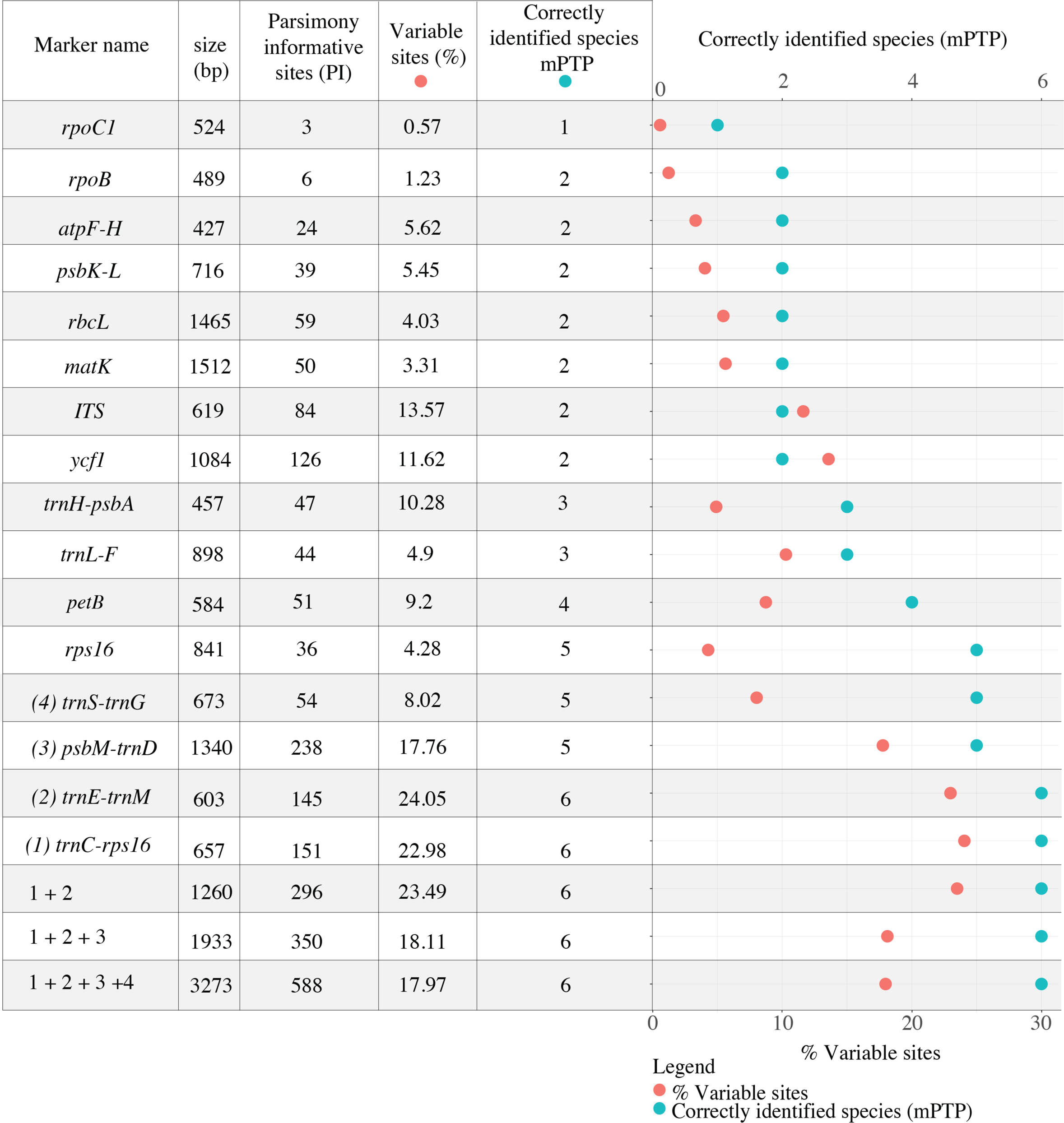
Percentage of variable site (orange) and successful species identified with the mPTP analyses (blue), for each marker and the concatenated matrices.

In the mPTP analysis for the full plastid dataset, the Average Support Value (ASV) assesses the congruence of support values with the ML delimitation. The analyses return an ASV of 97.9%, suggesting a high confidence for the given species delimitation scheme. Species delimitation recognized 21 distinct entities out of 20 species (Fig. S6). Over-representation and intraspecific variation of the *P. ginseng* samples has resulted in oversplitting this clade into two discrete entities. The *P. stipuleatus* / *P. binnatifidus* clade has lower data structure and the analyses does not strongly support the group as two independent mPTP entities (PP = 0.68). *P. quinquefolius* has been also divided into two subgroups, but the posterior probability of the subdivision is low (PP = 0.4).

The result of mPTP analyses for all previously used and the newly proposed markers are described in Figure 5 and the supported nodes for the speciation events have been added to the phylogenetic tree (supplementary material S7-S11). Out of the 15 analysed markers only four can be used to discriminate most species. Figure 5 also shows that regions with the highest density of parsimony informative sites are not necessarily the most efficient for species discrimination, and both skewed aggregated mutations as well as homoplasy can obscure phylogenetic patterns.

## Discussion

### Evolution of Araliaceae and ginsengs

The evolution of the Asian palmate group (Fig. 3) is concordant with previously published articles that show *Schefflera* at the base of the group. The paraphyletic genus *Dendropanax* was usually the most divergent in the group, but is now basal to the rest of the group. This position might be due to low sampling within the Asian palmate group. Results for *Brassaiopsis, Eleutherococcus, Fatsia, Kalopanax* and *Metapanax*, correspond with previously published phylogenies. Early radiations with interlineage hybridizations and genome doubling have been reported in the group [111] and this could explain the short internal branches. Further phylogenomic and biogeographical studies should be conducted to better understand the radiation of the Araliaceae.

In the Aralia-Panax group, Aralia is sister to *Panax*, and we find that *P. stipuleatus* forms a well-supported clade with *P. binnatifidus*, whereas previous studies have often reported that *P. binnatifidus* groups with *P. omeiensis, P. wangianus, P. zingiberensis* and *P. major* [19, 20, 38, 41], all four of which are however missing here. Due to the difficulty in obtaining material of *P. vietnamensis*, only three studies have included *P. vietnamensis* in a phylogeny [21, 96, 112]. The study by Lee et *al*. [112] using the plastid marker trnC–trnD does not resolve the position of *P. vietnamesis* in the phylogeny, but does identify a distinct clade consisting of *P. notoginseng, P. japonicus* and *P. vietnamensis*, which is also supported by our data. Komatsu et *al*. [21] recover a clade consisting of *P. vietnamensis* along with *P. japonicus* and *P. pseudoginseng* subsp. *himalaicus*, a synonym of *P. bipinnatifidus*. Inferring *P. japonicus* to belong to this clade is contradictory to previous studies that have found a clade consisting of *P. quinquefolius, P. ginseng* and *P. japonicus* [20, 38, 41, 112]. The plastome phylogeny supports a sister-relationship of *P. ginseng* and *P. quinquefolius*, the two economically most important species of ginseng. Although this full plastome phylogeny significantly differs from previously published molecular phylogenies, the new evolutionary pattern is strongly supported by bootstrap values and posterior probabilities.

### Incongruence between markers from different origin

Full length plastid genome data are a major improvement for the *Panax* phylogeny, and the addition of a bigger dataset has a strong influence on the phylogenetic hypothesis. However, discrepancies between full-length plastid genome phylogenies and nrDNA phylogenies are common in plants. nrDNA has been widely used for phylogenetic studies of *Panax* [19, 38, 41, 46], but the limitations of this approach have been extensively reviewed in [113]. Drawbacks of nrDNA include difficulties in aligning, and its limited use for phylogenetic inference between closely related and/or recently diverged taxa. It is also a challenge to determine the orthology and the paralogy of nrDNA sequences in the case of hybridization events or incomplete lineage sorting [114–116]. Bailey et *al*. [114] emphasise that despite valuable phylogenetic information from nrDNA, it might not the optimal choice to assess species trees, especially in case of allopolyploids or tetrapolypoids. Since this is also the case in *Panax*, we argue that nrDNA may be inappropriate to reconstruct the evolutionary history of this genus.

Phylogenetic congruence as well as incongruence of nuclear genomic and plastid marker data is well documented [117–119]. In the case of *Panax*, two of the nuclear markers used by [20] support the clade of *P. ginseng* and *P. quinquefolius* (Z14, Z8). However, our topology is incongruent for the remaining clades. Incongruences between the maternally inherited plastid genome and the biparentally inherited nuclear genes can be expected in genera with allopolyploid hybrids, like *Panax* [20]. Plastid phylogenies are not always representative of the species tree and might conflict with hypotheses of parsimonious morphological evolution [116, 120, 121]. Incongruences between plastome and nuclear gene trees have been reported in wide ranging groups of plants, such as *Asclepia* [72], *Helianthus* [122] and *Silene* [120].

### Enrichment

The novel method based on methylation-based enrichment increased the concentration of plastid DNA by 30% which is in the range found by a previous pilot study [84]. It is a suitable method for enriching the organellar genome before sequencing. The methylated fraction shows extremely low amounts of organellar DNA, meaning that we removed more than 99% of the non-methylated DNA from the total DNA. The *P. vietnamensis* sample had originally more degraded DNA and as a result shows a less successful enrichment. Using MBD2 to increase the concentration of organellar DNA in the total DNA allows multiplexing a larger number of samples. This method is appropriate for building plastid reference genome databases for barcoding projects. In case of degraded samples, we recommend removal of shorter DNA fragments before the enrichment.

### Selecting markers for molecular *Panax* identification

In DNA barcoding and plant product identification and authentication projects it is common to work with degraded DNA substrates for which it might be difficult to use methylation enrichment or the full plastid genome as a barcoding strategy. However, alternatives such as target enrichment and amplicon sequencing are possible [64, 123–125]. Here we have identified four variable regions that possess sufficient variation and genetic structure to discriminate most ginseng species. The identification of ginseng species is relatively complex because of the recent evolution and hybridization events. *P. ginseng* and *P quinquefolius* have recently diverged plastid genomes, and so do *P. binnatifidus* and *P. stipuleatus* [47]. Species delimitation using mPTP shows that for such species complexes traditional barcoding markers do not have enough structure for delimiting species. However, if carefully selected, some regions highlight specific structural patterns that enable the discrimination of species. The *trnC-rps16* region seems to be particularly promising, as it has enough variation to discriminate most species (Fig. S6). If plastid markers are to be used for barcoding, it is more relevant to use a combination of markers because mPTP analyses are better suited for multi-marker analyses [32]. A concatenated matrix with two, three or four markers combined improves the efficacy in segregating all the *Panax* species and specifically also those in closely related complexes. Our results suggest that a combination of the following markers: *trnC-rps16, trnE-trnM* and *psbM-trnD* (Fig. 5) enables confident identification of the main traded species *P. ginseng, P. quinquefolius* and *P. vietnamensis*. For further development, a complete sampling of all *Panax* species with multiple accessions per taxon should be made to confirm the observed variation in the selected markers.

In order to design accurate markers to monitor the trade of the medicinal species, it is necessary to understand the evolution of the targeted group. Many studies are based on the generic barcodes suggested by iBOL (*rbcL* and *matK*) without having strong evidence for the evolutionary hypotheses of the targeted group and a limited idea *a fortiori* of the discriminatory power of the used markers. Nonetheless, when a barcoding study targets a specific plant group or genus, and the barcode markers fail to yield a supported phylogeny, then one should aim to construct robust phylogenies with new markers to achieve species discrimination. If the phylogenetic hypothesis is not robust, or if the data are weak in structure as it is often the case with the standard barcoding markers, *rbcL* and *matK*, the resulting identifications might be misleading because of inaccurate species delimitation hypotheses [31].

The addition of genomic data for the phylogeny of *Panax* radically changes what is known about the evolution of the genus. The implications in terms of phylogeography are still unclear due to missing taxa, and the addition of population data and additional species should improve our insight into the evolutionary history of the genus. The development of species delimitation methods changes perspectives in molecular identification and DNA barcoding by incorporating evolution hypotheses at the species level. The newly proposed molecular markers allow for accurate identification of *Panax* species and enable authentication of ginseng and derived products and monitoring of the ginseng trade, while ultimately aiding conservation of wild ginseng.

## Declarations

### Acknowledgments

This project was supported by both the Vietnam Academy of Science and Technology (grant No. VAST02.01/16–17) and the European Union’s Seventh Framework Programme for research, technological development and demonstration under the Grant agreement no. 606895 to the FP7-MCA-ITN MedPlant, “Phylogenetic Exploration of Medicinal Plant Diversity”. The authors wish to thank the following people and organizations, Jarl Andreas Anmarkrud for his assistance for the enrichment, members of de Boer group for their helpful discussions and feedback during manuscript preparation. This work was performed on the Abel Cluster, owned by the University of Oslo and the Norwegian metacentre for High Performance Computing (NOTUR), and operated by the Department for Research Computing at USIT, the University of Oslo IT-department. http://www.hpc.uio.no/.

### Authors’ contributions

The project was coordinated by HLTT, VM and HdB. VM performed data analysis. VM and HdB wrote the main part of the paper. All other authors gave useful contribution on the analysis of data and text of the manuscript. All authors have read and approved the final version of the manuscript.

### Availability of supporting data

The raw sequence data from the *P. bipinnatifidus, P. stipuleanatus*, and *P. vietnamensis* samples have been to submitted to GenBank on the following accessions: SRR5725242, SRR5725240, SRR5725505, SRR5725492, SRR5738925, SRR5738922, SRR5738920, SRR5738920, SRR5738927.The DNA matrix used for the phylogenomic analyses are available on Open Science Framework, (doi: 10.17605/OSF.IO/Z7RWE). The plastome sequences of *P. bipinnatifidus, P. stipuleanatus*, and *P. vietnamensis* (2) have been submitted to NCBI GenBank, (MF377620, MF377621, MF377622, MF377623).

### Consent for publication

“Not applicable”

### Ethics approval and consent to participate

“Not applicable”

